# Predominance of animal-based expertise may bias NIH neuroscience grant review: A pilot study with implications for non-animal methodologies

**DOI:** 10.1101/2025.02.28.640877

**Authors:** Emily R. Trunnell, Katherine V. Roe

## Abstract

Despite increased enthusiasm for research methods that can replace the use of animals, the numbers of animals used in science and the funding for animal-based projects remain high. Our work addresses whether a possible “animal methods bias,” a preference for or greater familiarity with animal-based methods, in the grant review process may be contributing to this problem. In a novel pilot study to assess the possibility of animal methods bias in the National Institutes of Health’s (NIH’s) funding of basic, translational, and preclinical neuroscience research, we used the NIH’s iCite and RePORTER tools to characterize the expertise of members of 23 non-clinical NIH study sections and the methods used in successful grants scored by these review groups, based on publicly available data. Our data suggest that study sections assessing grants for basic, translational, and preclinical neuroscience research are largely composed of reviewers whose primary expertise is in animal-based methods. Animal use among reviewers positively correlated with the number of animal-based grants funded and negatively correlated with funding of non-animal research methods. Recommendations for further study and for avoiding or mitigating animal methods bias in grant review are provided.

## Introduction

In less than a decade, science has seen a global increase in the advocacy for—as well as the development and use—of new tools and systems that can replace the use of animals in biomedical research. Ethical and scientific arguments are equally at play. Public opposition to the use of animals in research is at an all-time high^1,2^ and, coupled with lower public trust in health officials and scientists and an interest in government spending habits,^3^ politicians are increasingly examining taxpayer investment in animal research and what technologies or changes to federal programs could reduce animal use. Since 2018, new programs that aim to reduce the use of animals in research and better support non-animal methods (NAMs, also called new approach methodologies) have been initiated in Australia, the E.U., India, the Netherlands, the U.K., and the U.S.^4–11^

In 2022, the NIH convened a working group to inform the agency’s director about where the use of NAMs could most benefit biomedical research. The working group issued its report in 2023 and the NIH director accepted its recommendations, which included prioritizing the development and use of NAMs, investing in NAMs training, and establishing and supporting infrastructure for NAMs tools.^12^ In recognizing the need for enhanced training in NAMs, the working group included grant reviewers as a target audience for this training “in terms of understanding proposals and the unique value of NAMs,” so that those who assess applications for funding may “appreciate the goal and technology behind the NAMs along with their usability for addressing specific research questions…[and] better understand how to evaluate the use of NAMs in fundamental and applied research grants.”^12^

NAMs include advanced *in vitro* models, such as organoids and organs-on-chips, *in silico* models that make use of computational techniques like machine learning to analyze human data and biological processes, and *in chemico* models to study how human molecules interact with each other and with external chemicals. Along with the potential to reduce animal use, NAMs can also provide information that more accurately translates to human health and disease, since the fundamental hurdle of species differences has been eliminated. To increase the use of NAMs in biomedical research, it is essential that individuals assessing grant applications have at least a basic technical understanding of these technologies and are able to accurately appreciate the potential for their use.

Because of the precedent set by the extensive history of animal use in biomedical research, many of the individuals charged with reviewing grants may have greater expertise and familiarity with animal-based methods. The preference for animal-based research methods or the lack of expertise to adequately evaluate NAMs has been termed “animal methods bias.”^13^ So far, this phenomenon has primarily been studied in the context of manuscript peer review, but survey and workshop proceedings have indicated its possible existence in areas other than publishing, such as funding, regulatory decision making, and science education.^14,15^

This analysis provides the first look into the potential for animal methods bias in grant funding, focusing on recent NIH review of basic, translational, and preclinical neuroscience research, an area with high translational failure from bench to bedside. Animal models for the study of neurodegenerative disease,^16,17^ psychiatric conditions,^18,19^ and other neurological disorders have been criticized for their lack of construct and predictive validity. With adequate funding, NAMs for neuroscience research may help improve basic and translational research, as well as new target discovery and preclinical testing in this field. In this pilot study, we use publicly available data to examine the potential for animal methods bias on the part of grant reviewers who compose NIH neuroscience study sections by assessing the technical expertise of these reviewers. We then compare reviewer expertise with the methods used in funded grants that were reviewed and scored by members of these study sections over a discrete period.

## Materials and methods

### Data collection: Review panel expertise

All data collection was performed between January 2023 and April 2024. The NIH’s Center for Scientific Review (CSR) website (https://public.csr.nih.gov/StudySections) provides publicly available descriptions of the research focus of review branches and study sections as well as information about group members. The 27 existing review branch descriptions at the time of data collection were evaluated to identify those whose role was to review neuroscience grant applications. Within those neuroscience-focused branches, study sections reviewing basic, translational, and preclinical neuroscience research, where proposed methods can include human, animal, or non-animal methods, were identified. Study sections that assess grant applications for clinical neuroscience research were excluded, as these are not expected to be replaced by NAMs. Membership rosters were downloaded from CSR’s public website and names of standing (chartered) members were extracted. Standing members only were included in this analysis for two reasons: 1) Standing members are said to play a larger role in grant review and have a greater influence on scoring than temporary members,^20^ and 2) Public records are not sufficiently transparent to allow for specific grants to be linked to specific review dates and their corresponding meeting rosters.

To calculate individual standing member’s methods expertise, individual names were entered into the NIH tool iCite (https://icite.od.nih.gov/analysis), which provides bibliometric data on the individuals’ peer-reviewed papers, including how ‘human-,’ ‘animal-,’ or ‘molecular/cellular biology’-oriented each paper is based on the fraction of MeSH terms included in the paper in each of these categories. Results were screened to ensure accuracy, comparing paper and journal titles to the individual’s field of work. Name disambiguation was performed when necessary, consulting institutional and professional websites to narrow the author pool by affiliations and middle initials. For individuals requiring name disambiguation, identifying characteristics were entered into a search on PubMed (https://pubmed.ncbi.nlm.nih.gov/) and resulting PMIDs were entered into iCite in lieu of the reviewer’s name.

For each of the standing members, the average human, animal, and molecular/cellular biology values from publications were extracted from iCite. These individuals’ scores were then used to calculate average expertise in these methods for each of the study sections included in the analyses (Table 1). iCite’s molecular/cellular category is not species-specific, including molecular methods that use both human and animal components, so this metric could not be used to assess reviewer expertise in NAMs research. Currently, no search tools exclusive and exhaustive for NAMs are available in the public domain.

**Table 1.**
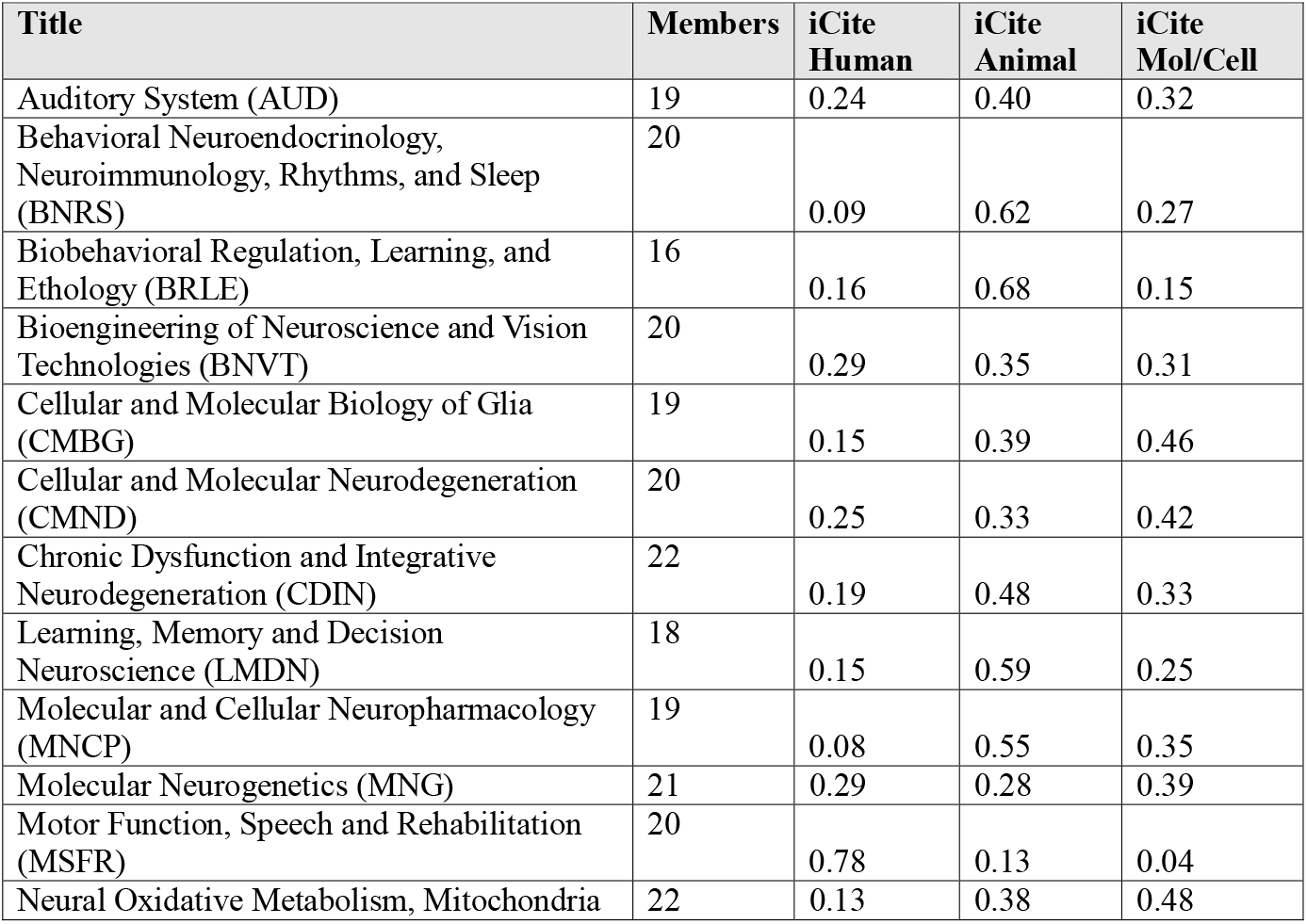
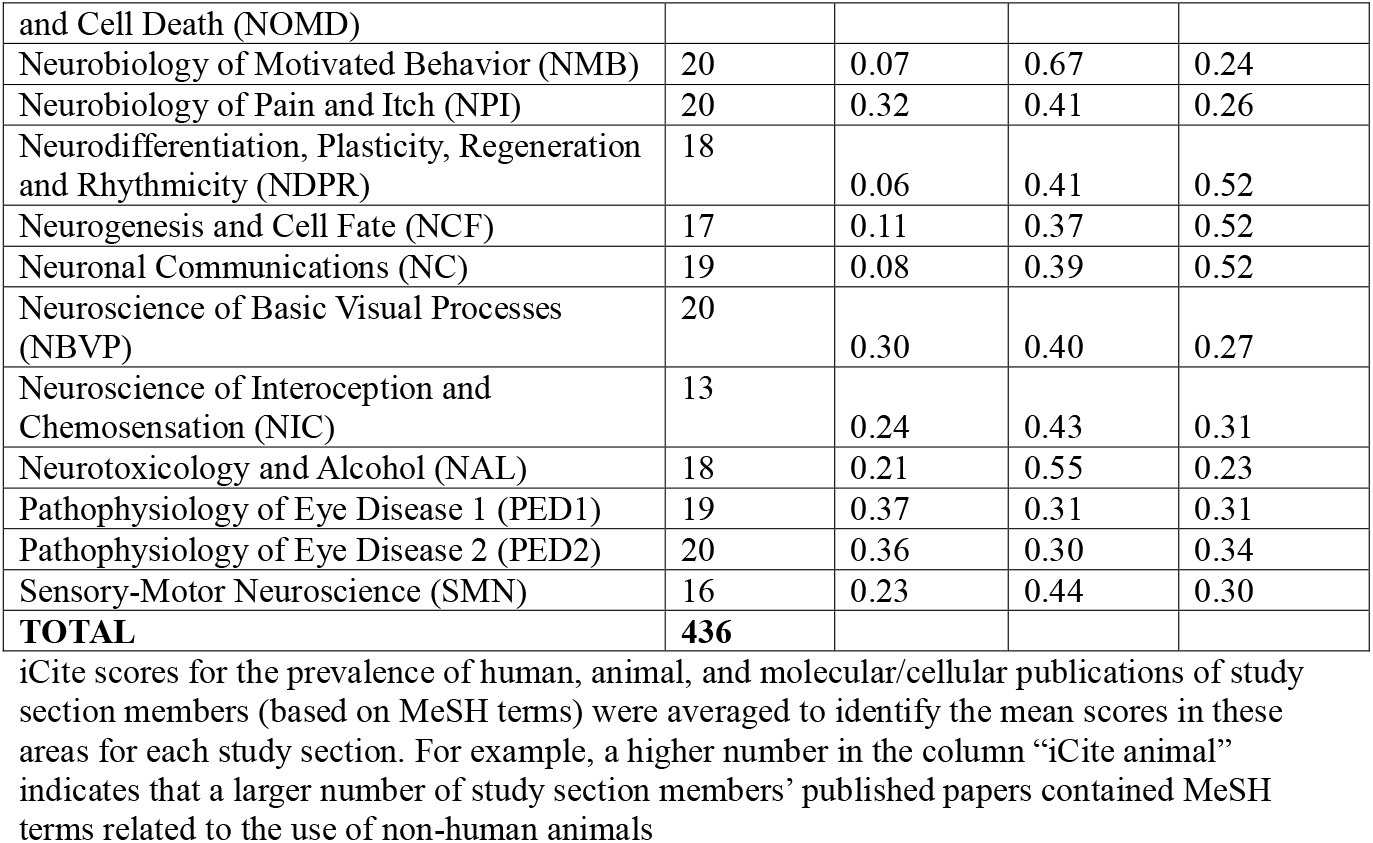
List of study sections assessing grant applications for basic, translational, and preclinical neuroscience research and the number of standing members on each study section.

### Data collection: Funding

Funded grants scored by each study section were retrieved from NIH RePORTER (https://reporter.nih.gov/), the agency’s electronic tool that allows users to search a repository of both intramural and extramural NIH-funded research projects and access publications and patents resulting from NIH funding. The search was focused to include only new grants (as new grants receive greater attention in study section meetings compared to grant renewals) and those awarded in the eight months prior to collection, as these were the grants most likely to have been reviewed by the current roster of study section members analyzed, based on posted meeting dates. Study section review typically occurs five months prior to the grant award, but our analysis was extended to eight months to account for possible delays. Project summaries were manually screened to determine whether the project proposed studies using humans, non-human animals or their cells or tissues, NAMs, or a combination of methods. When grant summaries were vague, methods sections of linked publications were referenced as well. Total project funding information was also extracted from RePORTER. For analysis of average grant costs per method, all grants using mixed methods were grouped into one category.

### Statistical analysis

All statistical analyses were performed using JASP (JASP Team (2024)); JASP (Version 0.18.3) statistical software. Average expertise in animal, human, or molecular/cellular was calculated for each of the 23 study sections included in this study. Similarly, the average use of different methods (animal, human, or NAMs) in all recent grants funded by the 23 study sections was also calculated.

To assess whether NIH study sections had differing levels of methods-type (animal, human, or molecular/cellular) expertise, and to determine whether funded grants differed in their use of method types (animal, human, NAMs), repeated measures Analysis of Variance (ANOVA) and post-hoc comparisons were performed. To determine the relationship between reviewers’ animal-methods expertise and the methods type used in subsequently funded grants projects, Pearsons’s correlation coefficients were calculated. The total amount of funding awarded by these study sections for grants using different research methods was also calculated. Raw data is publicly available in the OSF Data Repository at DOI 10.17605/OSF.IO/FKPZS.

### Ethics statement

No ethical approvals were required for this study. All data were obtained from publicly available databases.

## Results and discussion

### Study section methods expertise

Of the 27 existing NIH review branches at the time of data collection, we identified five whose role it was to review non-clinical neuroscience grant applications (Aging and Neurodegeneration, Basic Neuroscience, Biobehavioral Processes, Integrative and Cognitive Neuroscience, Neurotechnology and Vision). These review groups consisted of 38 study sections, 23 of which fit our inclusion criteria as being standing study sections charged with reviewing grants for basic, translational, and preclinical neuroscience research (Table 1). These 23 study sections were comprised of 436 standing members. Averaging individual members’ scholarly records, these study sections had significantly different expertise across the different methods types (F(2,44)=8.65, p<0.001), with an average of 43% animal, 22% human, and 33% molecular/cellular expertise across all 23 review panels (Table 1 and Fig 1). Post-hoc analyses indicated that the study sections had significantly more expertise in animal research methods than human research methods (t(22)=4.158, p<0.001). There was no significant difference in the study sections’ expertise in animal versus molecular/cellular methods (t(22)=1.993, p=0.157) or in their expertise in human versus molecular/cellular research methods (t(22)=2.165, p=0.107).

**Fig 1.**
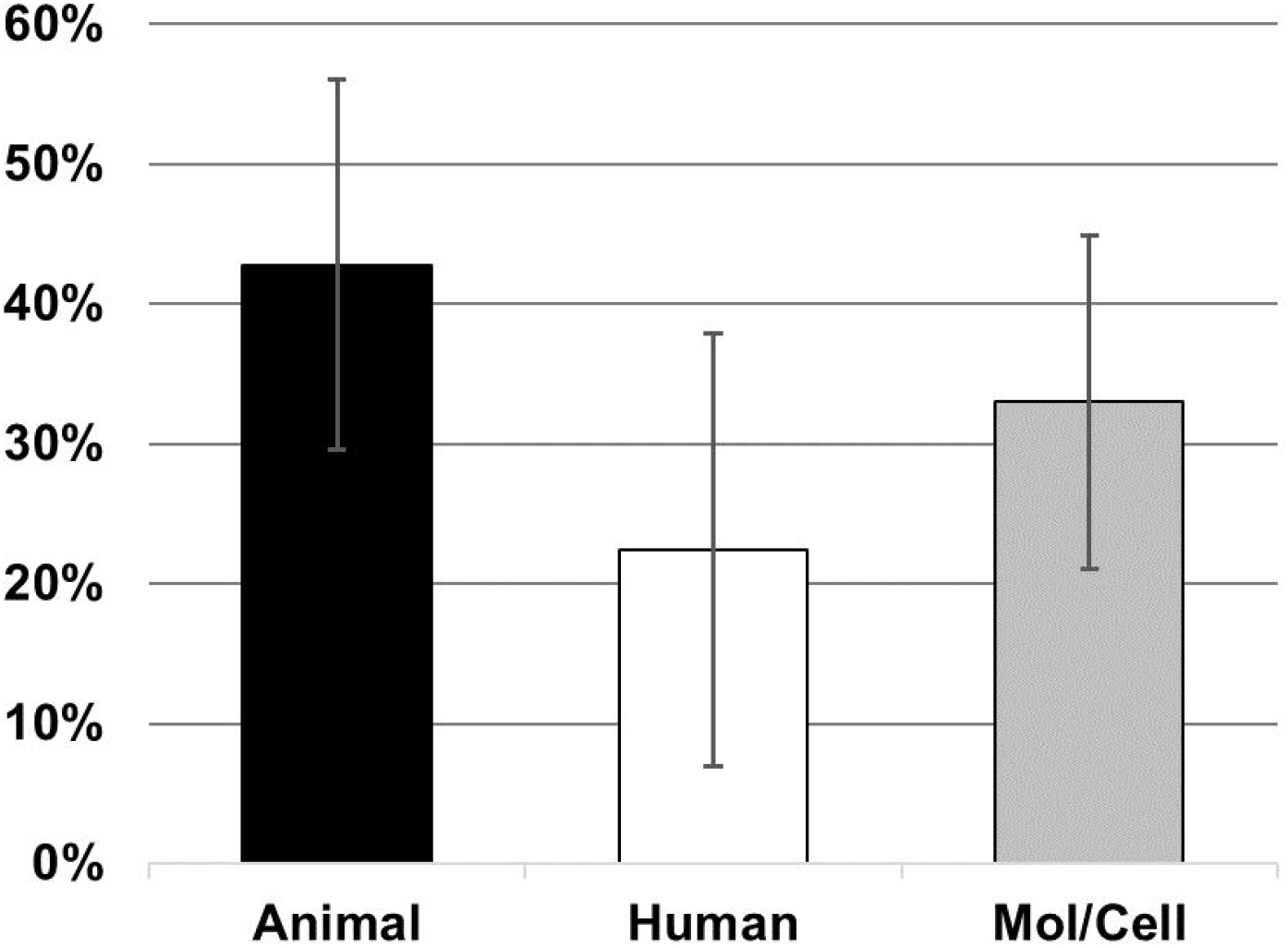
Average methods expertise of NIH basic, translational, and preclinical neuroscience study sections. Determined during the specified time frame by iCite analysis of relevant MeSH terms in each section members’ published papers. N=23; Error bars represent SEM.

### Research methods used in funded grants

Using NIH RePORTER, 679 funded grants were identified as having been scored by these 23 study sections in the eight months prior to data collection. For 117 grants (17%), the methods to be used were not discernible in the publicly available project summaries and primary research had not yet been published. Some grants proposed to use multiple methods. In these cases, each method was given equal weight in analysis. Of the 562 funded grants for which research methods were discernible, there was a significant difference in the type of methods used (F(2,44)=62.030, p < 0.001), with 72% proposing to use animals, 16% proposing to use NAMs, and 12% proposing to conduct human research (Fig 2). Post-hoc analyses indicated that the funded projects scored by these 23 study sections planned to rely significantly more on animal methods than either human methods (t(22)=9.978, p<0.001) or NAMs (t(22) = 9.276, p<0.001). There was no significant difference in funded grants’ proposed reliance on human versus NAM research methodology (t(22)=0.73, p=1.0).

**Fig 2.**
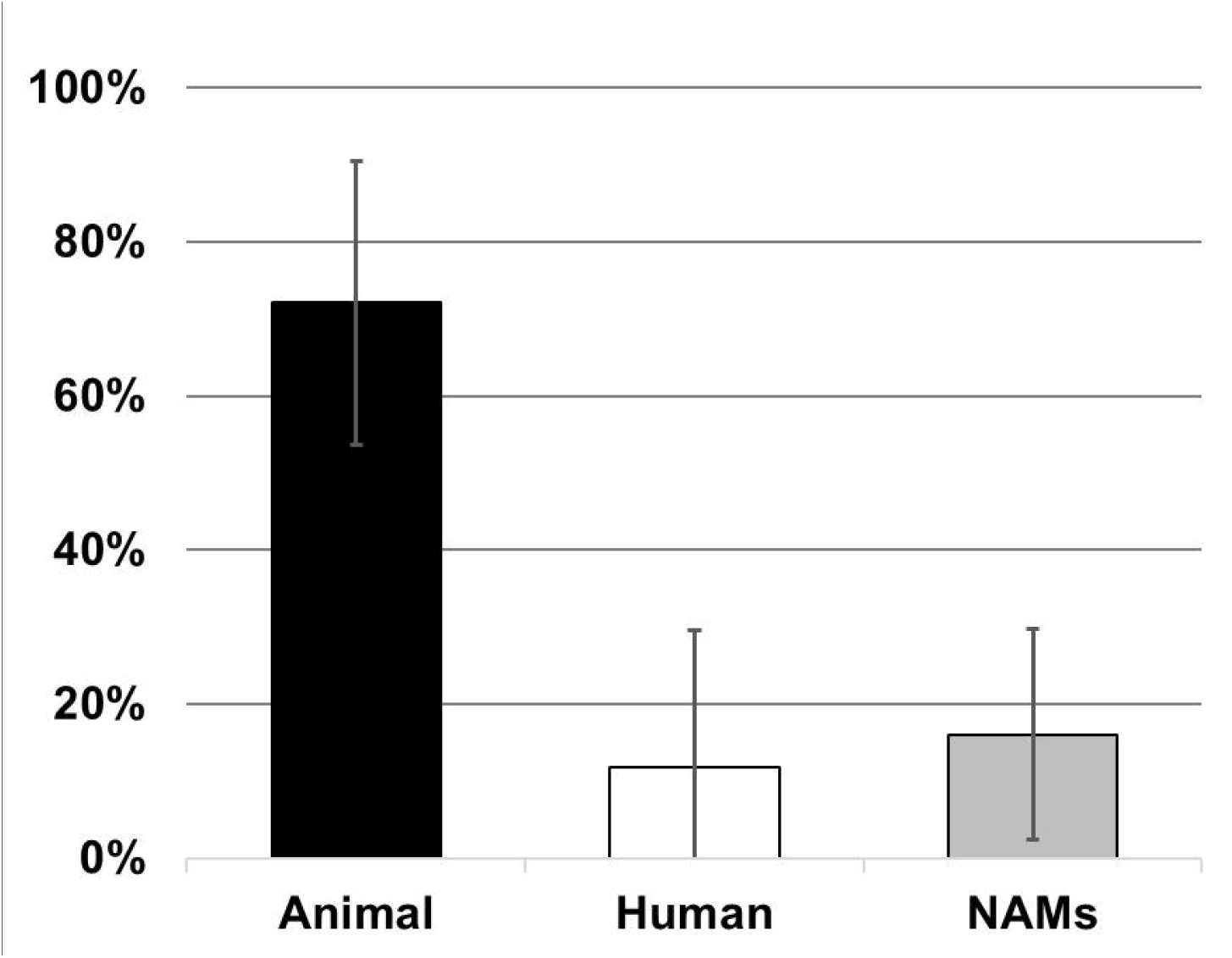
Percentage of funded grants using different method types scored by NIH basic, translational, and preclinical neuroscience study sections. N=562; Error bars represent SEM.

### Agency spending by research method

These 562 grants totaled $294,645,647 in federal funding. 454 (81%) of the grants proposed animal methods either alone or with other methods, accounting for $246,193,068 or 83.6% of agency funding for this type of research. 149 (26%) of grants used NAMs alone or in combination ($90,851,882; 30.8%) and 59 (10%) were for research on humans, either alone or combined with other methods ($25,968,431; 8.8%). The total amounts for funding of grants proposing single or mixed methods are shown in Fig 3. The high levels of funding for grants using only animal methods did not appear to be driven by differences in individual grant costs compared to grants using only humans or only NAMs, but costs associated with mixed-methods grants were higher than those using single methods (Fig 4). There was a significant main effect of methods-type on average grant costs (F(3, 558)=8.441, p<0.001), with mixed-methods grants receiving more money on average than animal-methods grants (t=4.319, p <0.001), human-methods grants (t=4.130, p<0.001), or NAMs methods grants (t=3.475, p=0.003). There was no significant difference in the average costs of single method grants.

**Fig 3.**
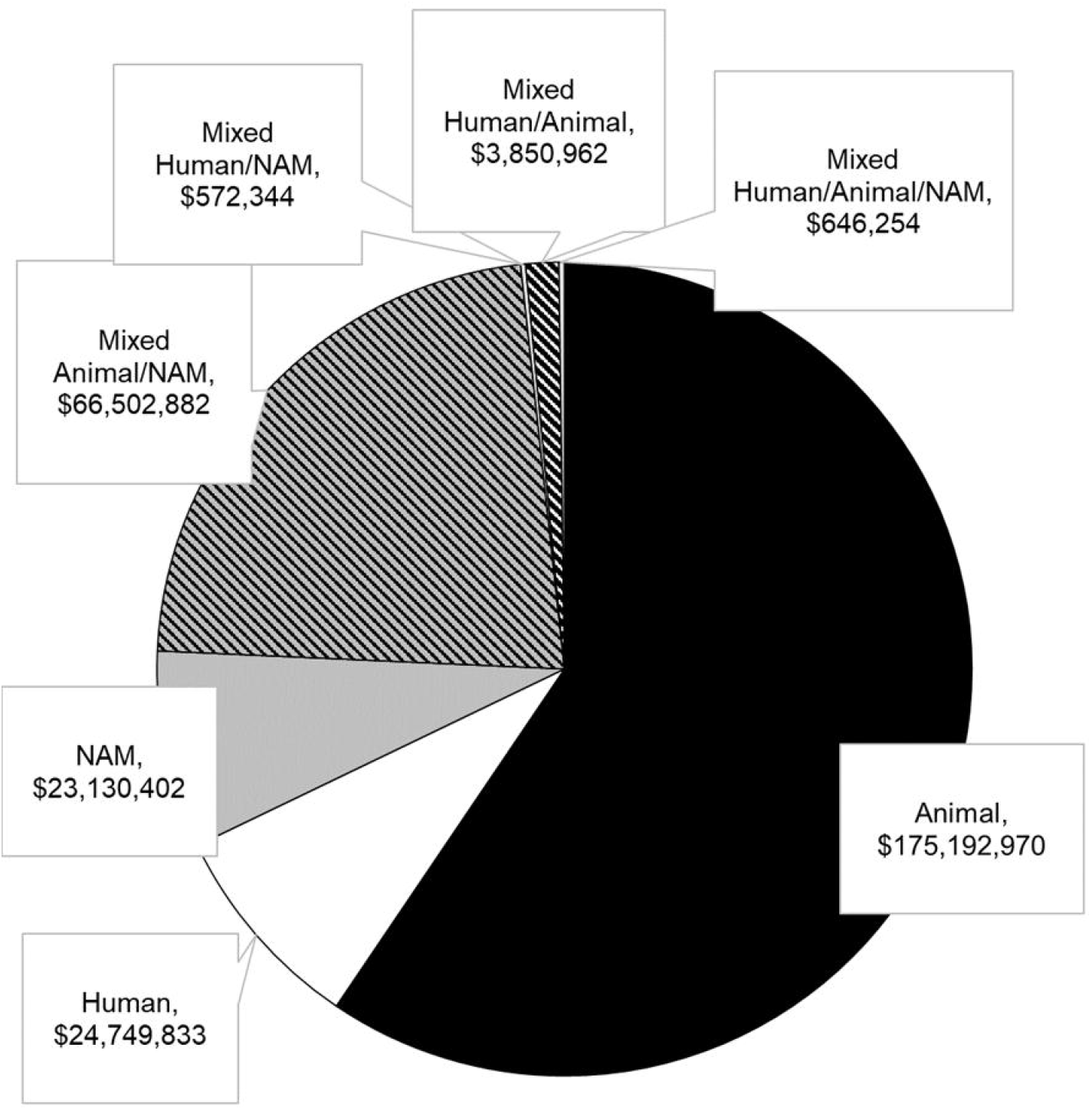
Total FY grant funding for projects proposing single or mixed methods which were scored by NIH basic, translational, and preclinical neuroscience study sections. N(Animal)=348; N(Human)=57; N(NAM)=50; N(Mixed Animal/NAM)=98; N(Mixed Human/NAM)=1; N(Mixed Human/Animal)=7; N(Mixed Human/Animal/NAM)=1; N(Total)=562.

**Fig 4.**
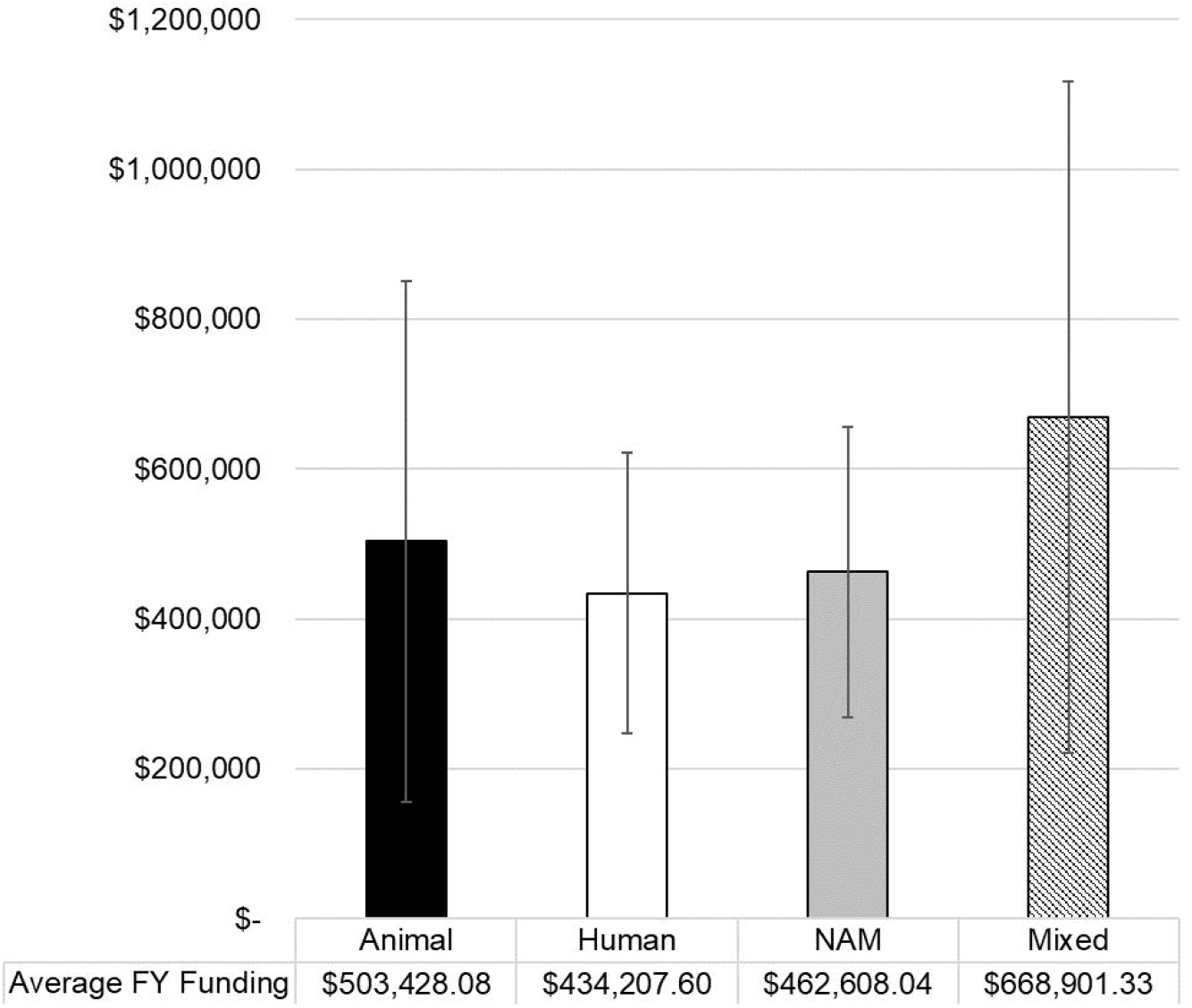
Average individual grant costs for NIH-funded basic, translational, and preclinical neuroscience projects. Includes grants proposing to use only animal (N=348), only human (N=57), only NAM (N=50) methods and those proposing to used mixed methods (N=107). Error bars represent SEM.

### Relationship between reviewer expertise and methods used in funded grants

Linear regression analysis revealed a strong positive correlation (r=0.769, p<0.001) between the study sections’ level of animal-based expertise and the reliance of animal-based methods in funded grants scored by those study sections (Fig 5a). There was a weak negative association (r=-0.365, p=0.0876) between animal-based expertise and the funding of grants using human methods (Fig 5b) and a moderate negative correlation (r=-0.568, p=0.005) between animal-based expertise and the funding of NAMs grants (Fig 5c).

**Fig 5.**
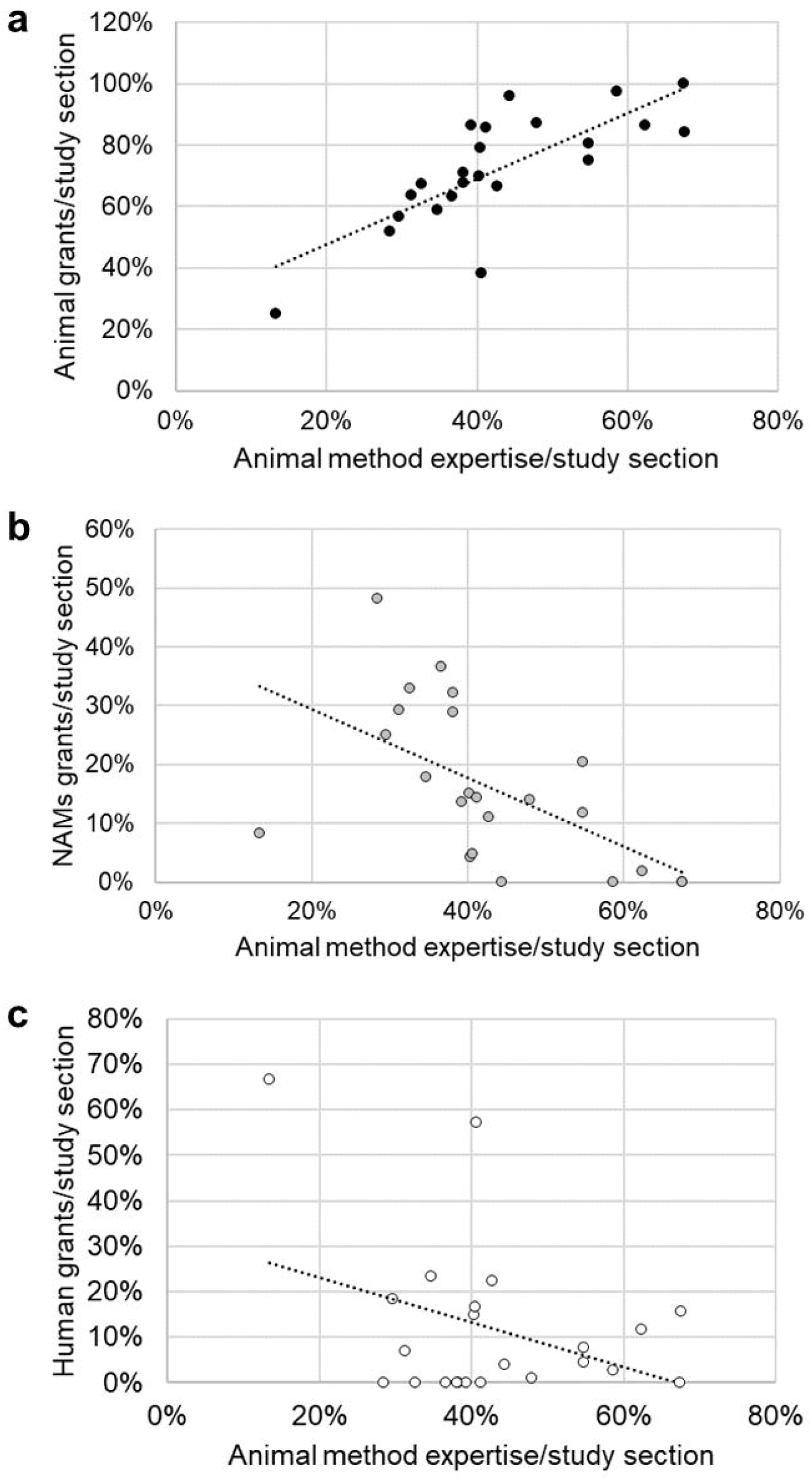
Linear regression analysis correlating animal method expertise per study section with methods used in grants funded by NIH basic, translational, and preclinical neuroscience study sections. a) R(animal expertise x animal grants)=0.769; b) R(animal expertise x NAMs grants)=-0.568; c) R(animal expertise x human grants)=-0.365.

## Discussion

Because of the long history of animal use in neuroscience, animal research became a discipline norm, embedded into institutional culture, and actors who uphold this culture may be more likely achieve success and be placed in positions of power, such as on peer review committees for grant applications or manuscript review.^21^ These individuals may, in turn, reinforce the culture of animal use either implicitly, perhaps through subconscious cognitive particularism^22^ or technical ignorance, or explicitly to quell the strong opposition to animal use that they perceive as a threat.^21^ The former could be partially mitigated by increased education and training in NAMs for existing grant reviewers and the early career researchers who will someday hold these positions.

These data indicated that the study sections tasked with reviewing NIH basic, translational, and preclinical neuroscience grant applications over an eight-month period were composed of individuals who predominantly held expertise in animal-based research methods and that funding for projects in this field, reflected both by the numbers of projects funded and the total funding allocated, chiefly supported animal-based methods. NIH funding for NAMs in basic, translational, and preclinical neuroscience was low: Only 26% of funded projects include NAMs and studies using only NAMs or NAMs in combination with human studies accounted for only 8% of agency funding for this type of research. Though larger and more comprehensive studies should be undertaken to understand this phenomenon in other research areas and over longer periods, this pilot study suggests that when study sections are populated by individuals with greater expertise in animal-based research methods, more funding is allocated towards animal-based research projects and less toward human-or NAMs-based research projects and that these differences are not driven by individual grant costs.

Data collection was limited, in part, by the inability to determine reviewer expertise in NAMs. Future analyses would benefit from the creation and curation of new search tools which are NAMs-specific and the integration of the tools into platforms like iCite. The U.S. National Toxicology Program has recently updated its ALTBIB search engine to allow users to query PubMed for topics related to specific uses or types of NAMs, for example, microphysiological systems.^23^ Aside from these specific cases, a formal search tool for all NAMs has yet to be developed.

A lack of transparency in project summaries of funded grants and information on unfunded projects were also limiting factors. Transparency continues to be an issue of interest for NIH and the agency has taken some steps as far as encouraging transparent reporting in grant applications and updates and for methods used in published, NIH-funded studies to aid reproducibility,^24^ but so far this initiative has not extended to the public-facing grant summaries available on RePORTER. For the funded grants in our dataset, 17% contained no information about the methods to be used in the study. Considering the public’s interest in science and how publicly supported research is conducted, NIH should mandate that project summaries describe whether studies are to be carried out using animals, humans, human specimens or data, a combination of these, or by other methods. Grant applications using human or animal subjects must already submit additional forms to the agency, so records for studies involving these populations already exist. Compiling information on grantees’ use of NAMs could also help the agency track its goals in better supporting these technologies.

Evidence of methods bias in NIH grant funding could be more rigorously assessed with the inclusion of information about grant applications that were not funded by the agency. Due to proprietary interests, information about unfunded grants is not released to the public. NIH’s Information for Management Planning Analysis and Coordination (IMPAC) II database is accessible to agency staff and could be used to provide a more complete picture about methods bias in funding or could be made available to meta-researchers studying this topic. Knowing whether NAMs grants are being rejected and having access to reviewer comments on NAMs applications would be the best indicator of whether reviewers have an adequate understanding of these systems and allow the agency to examine any evidence of animal methods bias. The recently published recommendations from the NIH Advisory Committee to the Director’s Working Group on Catalyzing the Development and Use of Novel Alternative Methods to Advance Biomedical Research acknowledged reviewers’ lack of awareness about NAMs, their potential advantages over animal-based methods, and how to properly evaluate them as a barrier to the broader development and use of these methods.^12^

These results provide initial data to support the need for reviewer training in NAMs and for reassessment of the balance of expertise on study sections. As this analysis is limited to neuroscience research and a discrete period, further studies should be done to assess whether the results apply to other fields of study and whether they hold over time.

## Conclusions

Despite public, political, and scientific support for transitioning away from the use of animals in science, funding for methods that can replace the use of animals remains low. The predominance of animal-based methods expertise among grant reviewers in neuroscience over the period assessed in this pilot study appears to correlate with high funding for animal-based research projects and low funding for projects which use NAMs.

To ensure that animal methods bias is not unfairly affecting the funding of projects proposing to use NAMs and thus serving as a barrier to their uptake, the NIH and other research funders should:

- Conduct internal studies assessing animal methods bias frequency and impact within grant review
- Broaden the pool of NAMs expertise available in reviewer pools
- Create NAMs-specific funding streams
- Encourage applicants to report incidences of animal methods bias during grant review
- Consult the Coalition to Illuminate and Address Animal Methods Bias (COLAAB, www.animalmethodsbias.org), an international collaboration of researchers and advocates (of which the corresponding author is a member), as it develops strategies to assess and mitigate animal methods bias in grant funding
- Foster transparency and meta-research by ensuring that publicly available grant summaries include the methods to be used in the research
- Facilitate meta-research on unfunded grants

It is critical that funding agencies assess barriers to the wider adoption of NAMs in order for the U.S. to support a research workforce that is in line with the global movement towards the reduction of animal use in biomedical research. Facilitating this transition to NAMs has the potential to improve the human translatability of federally funded research and to meet societal expectations.

## Acknowledgements

The authors would like to thank Kati Bertrand for her assistance in data collection.

## Notes

https://osf.io/fkpzs/?view_only=6fdc6b4c2edd438e9d11af0437466337

### Competing Interest Statement

The authors have declared no competing interest.

